# Evolution, not transgenerational plasticity, explains the divergence of acorn ant thermal tolerance across an urban-rural temperature cline

**DOI:** 10.1101/571521

**Authors:** Ryan A. Martin, Lacy D. Chick, Aaron R. Yilmaz, Sarah E. Diamond

**Affiliations:** Department of Biology, Case Western Reserve University, Cleveland, Ohio 44106

**Keywords:** adaptation, maternal effect, thermal physiology, global change, speciation

## Abstract

Disentangling the mechanisms of phenotypic shifts in response to environmental change is critical, and although studies increasingly disentangle phenotypic plasticity from evolutionary change, few explore the potential role for transgenerational plasticity in this context. Here, we evaluate the potential role that transgenerational plasticity plays in phenotypic divergence of acorn ants in response to urbanization. F2 generation worker ants (offspring of lab-born queens) exhibited similar divergence among urban and rural populations as F1 generation worker ants (offspring of field-born queens) indicating that evolutionary differentiation rather than transgenerational plasticity was responsible for shifts towards higher heat tolerance and diminished cold tolerance in urban acorn ants. Hybrid matings between urban and rural populations provided further insight into the genetic architecture of thermal adaptation. Heat tolerance of hybrids more resembled the urban-urban pure type, whereas cold tolerance of hybrids more resembled the rural-rural pure type. As a consequence, thermal tolerance traits in this system appear to be influenced by dominance rather than being purely additive traits, and heat and cold tolerance might be determined by separate genes. Though transgenerational plasticity does not explain divergence of acorn ant thermal tolerance, its role in divergence of other traits and across other urbanization gradients merits further study.

## Introduction

Responses to changing environments can occur either through evolutionary change or through existing phenotypic plasticity (Diamond and Martin 2016). However, the mechanisms underlying observed phenotypic responses to rapid environmental changes, including through anthropogenic effects are largely unknown for most organisms, as distinguishing plastic from evolved responses is often challenging (Merilä and Hendry 2014). Indeed, while the dichotomy between purely plastic and purely evolutionary responses is important and real, the two mechanisms can interact and influence each other (Ghalambor et al. 2007; Diamond and Martin 2016; Kelly 2019). This more complex view posits that along with the concept that plasticity itself evolves (DeWitt and Scheiner 2004), plastic responses to novel environments may often precede and shape the speed, direction, and magnitude of evolutionary change, thereby facilitating adaptation (*i.e., plasticity-first hypothesis* or *genetic accommodation*; for theoretical considerations, see Price et al. 2003; West-Eberhard 2003; Levis and Pfennig 2016, and for empirical examples, see Corl et al. 2018; Levis et al. 2018). Transgenerational effects—the influence of parents on the phenotype of their offspring independent from the effects of genetic inheritance (Bernardo 1996; Mousseau and Fox 1998; Marshall and Uller 2007)—may be especially effective in facilitating adaptive evolution to novel environments, as parents can alter offspring development in anticipation of the environment they will later experience (Badyaev 2008). Despite this promise, there are few examples of transgenerational plasticity facilitating evolution in response to the altered selective regimes of novel environments (Badyaev et al. 2002a,b, Pfennig and Martin 2009, 2010).

Cities are an emerging experimental model system for studying plastic and evolutionary responses to rapid anthropogenic change (Rivkin et al. 2019), and a number of studies find evidence that populations are indeed evolving in response to selective pressures associated with urbanization over contemporary timescales (Johnson and Munshi-South 2017). For example, research suggests that urban populations of the Puerto Rican Crested Anole, (*Anolis cristatellu*s) have evolved longer limbs and increased toe pad lamellae in response to selection for locomotor performance on artificial surfaces (Winchell et al. 2016). This and other studies have also found considerable within generational phenotypic plasticity in response to urban environments (*e.g.*, Diamond et al. 2017, 2018a; Gorton et al. 2018). However, evaluating if transgenerational plasticity contributes to divergence in urban habitats has not yet been explored.

A classic approach for disentangling plastic responses occurring both within and across generations from evolved responses is to compare field phenotypic measures between populations with those from individuals of the same populations bred and reared in a controlled, shared environment (*i.e.*, a common garden experiment) across multiple generations (Merilä and Hendry 2014; Donihue and Lambert 2015). By controlling for environmental differences across populations in the common garden, researchers can isolate the genetic component, if any, of the phenotypic divergence measured in the field. Although this approach is conceptually simple, such experiments are often labor intensive and time consuming, and might be challenging or impossible for species that are difficult to rear and breed in the lab (Donelson et al. 2018). Consequently, most urban evolution studies to date have only reared field-caught juveniles or a single generation in the lab and as a result cannot fully disentangle transgenerational phenotypic plasticity from within generation plasticity and purely genetic divergence (but see Brans et al. 2017).

Here we use a laboratory common garden experiment with a multi-generational breeding design to test for the presence of transgenerational plasticity and also investigate the genetic architecture underlying evolutionary divergence between urban and rural populations of the acorn ant (*Temnothorax curvispinosus*). We focus on thermal tolerance traits (heat and cold tolerance) and their response to urban heat island effects along the urbanization gradient of Cleveland Ohio, USA. F1 offspring (lab-born workers of field-caught queens) of urban acorn ant populations reared under common garden exhibit increased heat tolerance and diminished cold tolerance compared with rural populations; this pattern of differentiation among urban and rural acorn ant populations is apparent for geographically isolated cities, including Cleveland, Ohio and Knoxville, Tennessee (Diamond et al. 2017, 2018a). Moreover, in both cities we found that urban colonies produced more alates (*i.e.*, reproductive offspring) under typical urban rearing temperatures, and rural colonies produced more alates when reared at typical rural rearing temperatures, resulting in fitness tradeoffs across environments for each population (Diamond et al. 2018a). These results strongly suggest that *T. curvispinosus* has adaptively evolved to the warmer environmental temperatures of the urban environment (*i.e.*, the urban heat island effect). However our design could not detect or rule out transgenerational plasticity as a mechanism behind this adaptive divergence.

In this study, we tested the heat and cold tolerance of F2 offspring, lab-born workers of lab-born queens. We tested the offspring of pure-type rural population matings, pure-type urban population matings, and hybrids, both where the female was from the rural population and the male from the urban population and vice versa. If transgenerational plasticity were responsible for the increased heat tolerance and diminished cold tolerance we documented previously in urban population acorn ants, then we would expect these phenotypic differences to disappear in the F2 offspring comparisons of pure-type urban and rural populations. If transgenerational plasticity plays only a minor to no role, we would expect the increased heat tolerance and diminished cold tolerance of pure-type urban populations relative to rural populations to remain in the F2 offspring. The hybrid matings further evaluate the role of transgenerational plasticity, where parental effects should differentially affect hybrid offspring depending on the direction of the urban-rural mating. Additionally, comparisons among and against the hybrid tolerance phenotypes allowed us to gain further insight into the genetic architecture underlying heat and cold tolerance. For example, these comparisons allowed us to distinguish a purely additive genetic model of trait inheritance from alternatives, such as dominance or transgressive segregation.

## Materials and Methods

### Colony collections

We collected queenright (queen present) acorn ant (*Temnothorax curvispinosus*) colonies in early summer 2017 (6 June 2017 – 21 July 2017) from urban and rural sites across Cleveland, Ohio (42°N latitude). To identify urban and rural sites, we used percent developed impervious surface area (ISA) (Imhoff et al. 2010), such that urban sites were categorized as 40-50% ISA, and rural sites were categorized as 0% ISA (see Table S1 for site details).

### Mating design and common garden rearing

We returned field-caught colonies to the lab and placed them under conditions to facilitate the production of male and female alates (or winged, sexual individuals), including a warm, diurnally fluctuating thermal regime (30 °C daytime, and 26 °C nighttime temperature) synced with a 14:10 short-day photoperiod (following Stuart et al. 1993; Percival 36-VL growth chambers). Each colony was maintained individually in a 120 mL plastic cup. Resource tubes with sugar water (25% solution) and plain tap water were provided to colonies along with a continuous supply of dead mealworms. Once alates were produced within the colonies, we paired males and females in four mating crosses, including two pure-type matings, rural-rural and urban-urban and two hybrid matings, rural-urban (with the maternal source population listed first) and urban-rural. Because acorn ants are suggested to have male-biased dispersal (Stuart et al. 1993), we provided several female alates (typically 5 alates) from the same colony with males from several different colonies (typically 10 alates, all of which were distinct from the female alate source colony). We enclosed alates from each cross in separate glass aquaria (38 L capacity) with a mesh netting top. Alates were provided with the same resources as their source colonies, including water and sugar tubes plus dead mealworms. We provided alates with several uninhabited, hollowed-out acorns that we had sliced horizontally and held together with garden twine. This allowed us to check on the mating status of the paired alates, specifically whether the ants had shed their wings (an indication that they have mated, (Herbers 1990) and whether mated female alates began to lay eggs. All mated females began new egg production by 7 December 2017.

Once mated female alates began to produce eggs, we placed these newly established colonies individually within 120 mL plastic cups and provided them with water and sugar resource tubes and dead mealworms. We held these colonies at 25 °C (on a 14:10 photoperiod), as this is the optimal temperature for brood development (Diamond et al. 2013). After the newly laid eggs developed into workers, we tested the thermal tolerance of these workers (*i.e.*, the second generation of acorn ant workers reared entirely within the laboratory environment).

### Thermal tolerance

We used a dynamic temperature ramping protocol to assess the critical thermal maximum and minimum (CT_max_ and CT_min_), each defined as the loss of muscular coordination, which yield ecologically relevant limits on performance and serve as our measures of heat and cold tolerance (Terblanche et al. 2011). We tested workers individually for either CT_max_ or CT_min_, as the assessment of thermal tolerance is a semi-destructive process that precludes assessment of both CT_max_ and CT_min_ on the same individual. Ants were placed individually into 1.5 mL Eppendorf tubes with a cotton plug in the lid. Temperatures were manipulated using a dry block incubator (Boekel Scientific Tropicooler), and increased or decreased at a rate of 1 °C min^−1^. Initial temperature for the estimation of CT_max_ was 34 °C, and was 16 °C for CT_min_. These starting temperatures lie outside the range needed to induce loss of muscular coordination or death.

Thermal tolerances were assessed between 30 August 2018 and 2 September 2018. We tested a total of 487 individuals for thermal tolerance, 251 CT_max_, 236 CT_min_ from 18 different colonies. For the pure mating types, we had four urban-urban colonies and four rural-rural colonies. For the hybrid mating types, we had five urban-rural (with the origin of the maternal population listed first) colonies and five rural-urban colonies. The number of individuals tested per colony (and within each tolerance type, heat or cold tolerance) ranged from 5 to 24 with a mean and SD of 13.6 and 5.17 (see also Table S2).

### Statistical analyses

We fit linear mixed effects models with either CT_max_ or CT_min_ as the response variable and the type of mating (the two pure types, urban-urban and rural-rural, and the two hybrid types with the population origin of the mother listed first, urban-rural and rural-urban) as a predictor variable (treated as a categorical variable with four levels). We used the *lme* function from the *nlme* library in R (Pinheiro et al. 2018; R Core Team 2018) to perform these models. Colony identity was included as a random intercept in each model to account for autocorrelation among individuals from the same colony. The statistical significant of mating type was assessed using likelihood ratio tests. Post-hoc analyses to assess the significant differences between factor levels of the type of mating predictor variable were performed using the *emmeans* function (from the eponymous library) with all pairwise comparisons (Lenth 2018).

To model tolerance breadth (the difference between colony-mean CT_max_ and colony-mean CT_min_), we fit a generalized linear model with tolerance breadth as the response and type of mating as a predictor. We modeled tolerance breadth using a Gaussian probability distribution and log link function owing to skew in the untransformed data. Because hybrid tolerance responses were indistinguishable from one another, and because we were more limited in statistical power after computing colony-level means for heat and cold tolerance, we fit a subsequent model wherein each of the pure types (urban-urban and rural-rural) were compared with the hybrid types (considered as a single group). We performed the post-hoc analysis on this model. Although a strict test of maternal effects would involve comparisons of the urban maternal pairings versus rural pairings, our initial analyses showed little evidence of maternal effects (see below), so we used the particular analyses for thermal tolerance breadth to explore the genetic architecture underlying heat versus cold tolerance, and hence justified our combining of offspring results for the hybrid matings.

## Results

We found no evidence that the evolutionary differentiation in heat and cold tolerance among urban and rural acorn ant populations was driven by transgenerational plasticity. Likelihood ratio tests for the effect of mating type on heat tolerance and cold tolerance were statistically significant (CT_max_: *χ2* = 44.6; *P* = 1.12E-09; *df* = 3; CT_min_: *χ2* = 20; *P* = 0.00017; *df* = 3). Post-hoc tests revealed that F2 offspring of male and female reproductives born, reared, and mated within the lab environment exhibited significantly elevated heat tolerance and significantly diminished cold tolerance in urban-urban crosses as compared with rural-rural crosses (Table 1, Fig. 1).

**Table 1.** Post-hoc analyses of the mating type term for models of CT_max_, CT_min_, and tolerance breadth are provided including estimates of the contrast, their standard errors and test statistics and *p*-values for the pairwise differences between factor levels of the mating type term. Note that hybrid factor levels are combined for the analysis of tolerance breadth, and that estimates and test statistics are reported on the natural log scale. Significant *p-*values at the 0.05 level are indicated in bold font.

| Tolerance type | Contrast | Estimate | SE | Test statistic | *P* |
| --- | --- | --- | --- | --- | --- |
| $CT_{\max}$ | rur-rur - rur-urb | -1.25 | 0.257 | -4.84 | **0.00132** |
|  | rur-rur - urb-rur | -1.33 | 0.257 | -5.17 | **0.000723** |
|  | rur-rur - urb-urb | -1.67 | 0.272 | -6.17 | **0.000128** |
|  | rur-urb - urb-rur | -0.0855 | 0.245 | -0.35 | 0.985 |
|  | rur-urb - urb-urb | -0.428 | 0.259 | -1.65 | 0.384 |
|  | urb-rur - urb-urb | -0.343 | 0.26 | -1.32 | 0.566 |
| $CT_{\min}$ | rur-rur - rur-urb | -0.304 | 0.405 | -0.75 | 0.875 |
|  | rur-rur - urb-rur | -0.331 | 0.408 | -0.813 | 0.848 |
|  | rur-rur - urb-urb | -1.75 | 0.43 | -4.08 | **0.00548** |
|  | rur-urb - urb-rur | -0.0276 | 0.39 | -0.0709 | 1 |
|  | rur-urb - urb-urb | -1.45 | 0.412 | -3.51 | **0.0161** |
|  | urb-rur - urb-urb | -1.42 | 0.415 | -3.42 | **0.019** |
| Tolerance breadth | hybrid - rur-rur | 0.0231 | 0.00984 | 2.35 | **0.0490** |
|  | hybrid - urb-urb | 0.0260 | 0.00986 | 2.64 | **0.0225** |
|  | rur-rur - urb-urb | 0.00290 | 0.0119 | 0.244 | 0.968 |

**Figure 1.**
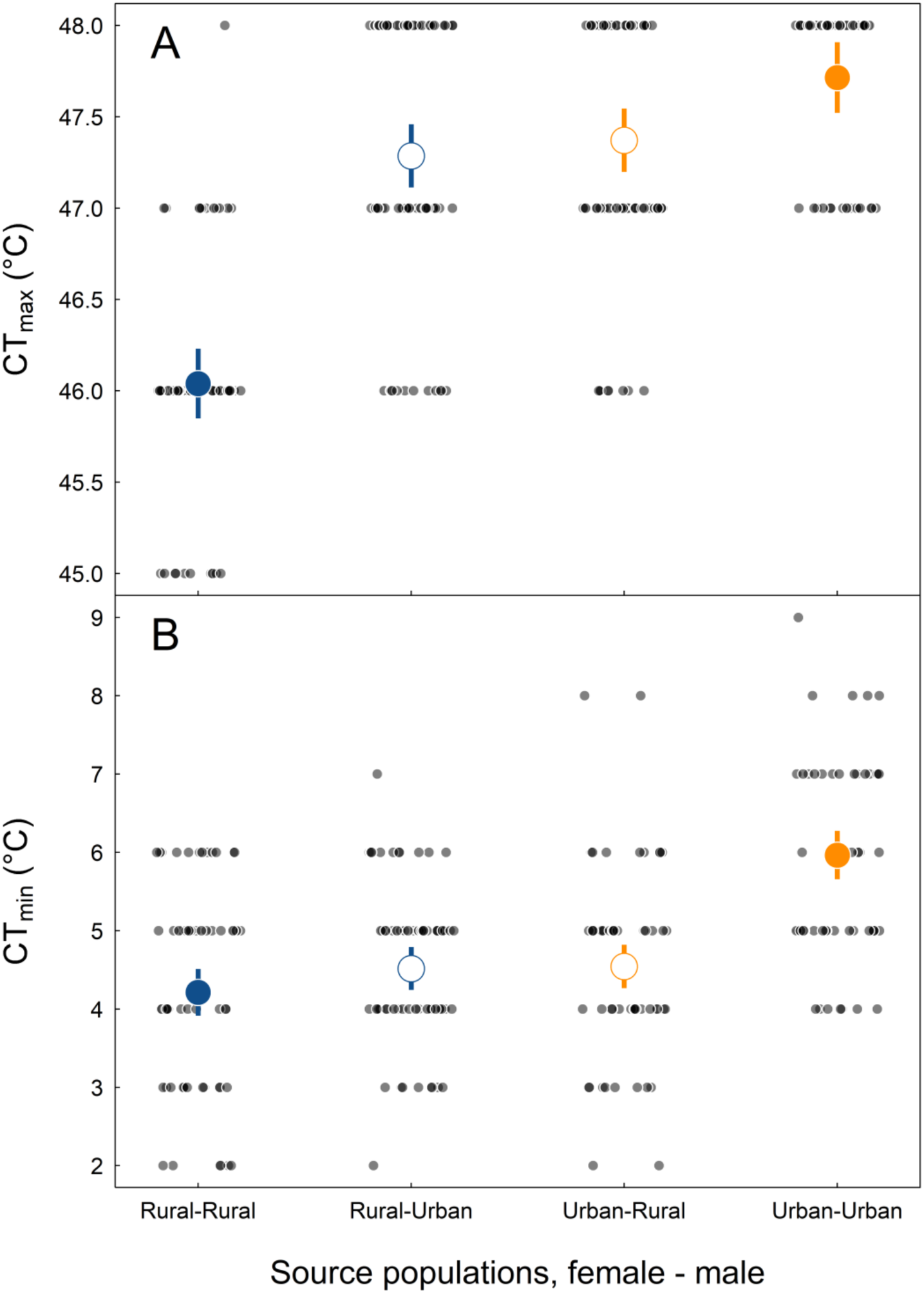
(A) Heat tolerance (CT_max_, °C) and (B) cold tolerance (CT_min_, °C) as a function of pure type and hybrid matings across urban and rural acorn ant populations. Small points indicate tolerance values of individual worker ants from each mating type (with the maternal source population listed first): rural-rural, rural-urban, urban-rural, and urban-urban. Predicted mean heat tolerances and standard errors from a linear mixed effects model that accounts for colony-level autocorrelation are presented in large points (means) and line segments (standard errors). Pure types are in filled symbols (blue for rural and orange for urban). Open symbols represent hybrid matings, with the color reflecting the maternal population origin.

The results of the hybrid pairings (female-male) urban-rural and rural-urban revealed further insights into the contemporary evolution of thermal tolerance traits in response to urbanization. First, the fact that the hybrid matings were successful in producing offspring indicates no evidence of behavioral prezygotic or intrinsic postzygotic reproductive isolation among urban and rural populations. It is possible that F2 offspring may exhibit later signs of reproductive isolation (hybrid inviability or sterility). We also note that male F2 hybrid incompatibility is unclear from our experimental design and study system as males are haploid clones of diploid queens, and thus another round of mating (F3) would be required to test this possibility (Koevoets and Beukeboom 2009). In any case, these are secondary goals of our study.

Interestingly, hybrids were more similar to either the rural-rural or urban-urban pure types depending on whether heat tolerance or cold tolerance was assessed. The heat tolerance of hybrids more resembled the urban-urban pure type than the rural-rural pure type (Fig. 1A), whereas the cold tolerance of hybrids more resembled the rural-rural pure type than the urban-urban pure type (Fig. 1B). This provides further evidence that divergence in thermal tolerance does not stem from transgenerational plasticity, as the offspring did not consistently resemble a specific parental population. Moreover, this differential matching of hybrids and pure types across heat and cold tolerance potentially suggests some degree of genetic decoupling of the traits.

We further explored the potential ecological consequences of the differential matching of hybrid and pure-type heat and cold tolerance phenotypes by computing their tolerance breadth (difference between colony mean CT_max_ and CT_min_). The likelihood ratio test for the effect of mating type on tolerance breadth was statistically significant (*χ2* = 9.22; *P* = 0.0266; *df* = 2). Post-hoc tests revealed that hybrids exhibited greater tolerance breadth than either of the pure types since the hybrids possessed both the superior heat tolerance value (high CT_max_) of urban populations and the superior cold tolerance value (low CT_min_) of rural populations (Figure 2; Table 1).

**Figure 2.**
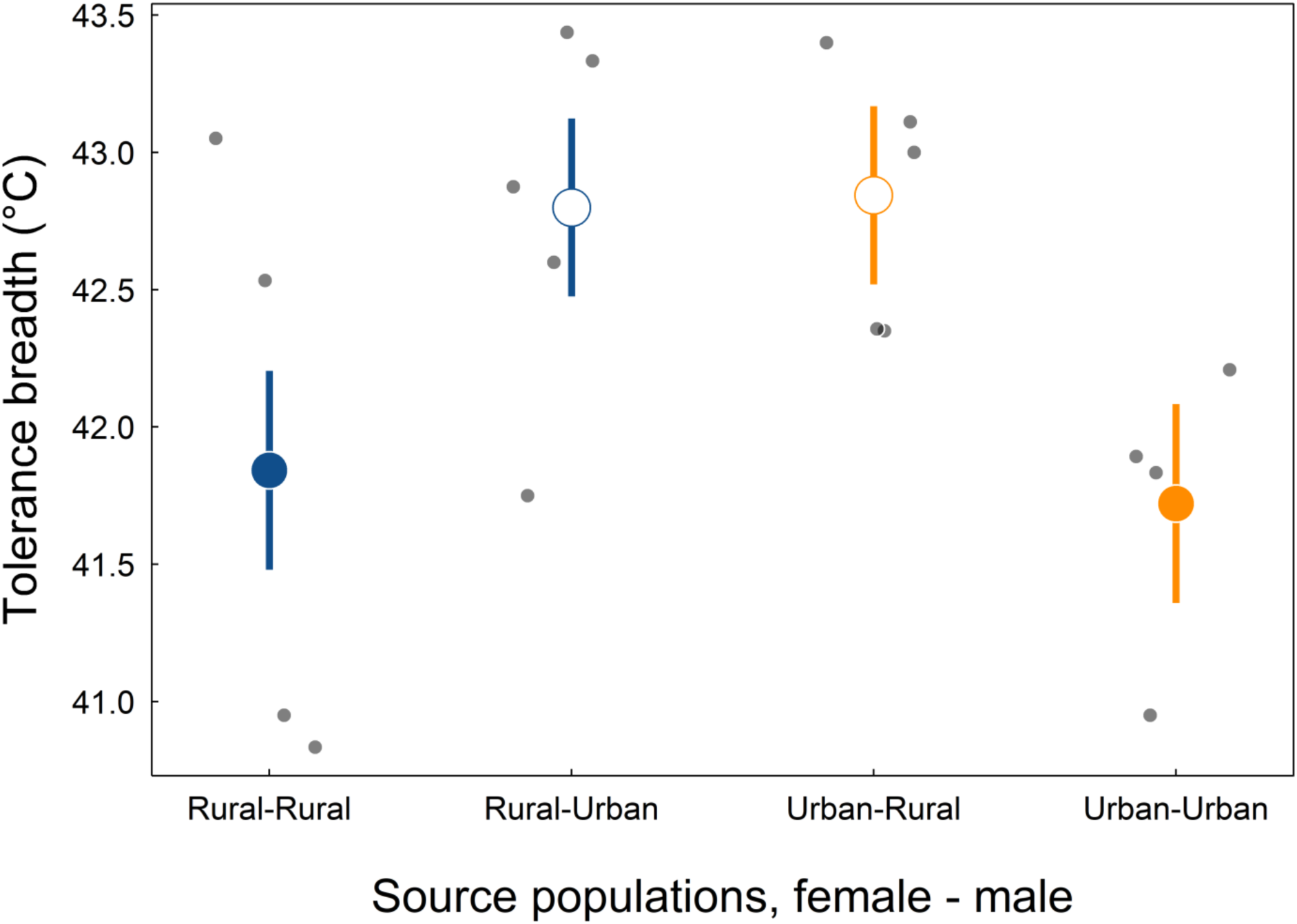
Tolerance breadth (°C) as a function of pure type and hybrid matings across urban and rural acorn ant populations. Small points indicate tolerance breadth values of whole colonies (colony mean CT_max_ - colony mean CT_min_) from each mating type (with the maternal source population listed first): rural-rural, rural-urban, urban-rural, and urban-urban. Predicted mean tolerance breadths and standard errors from a generalized linear model are presented in large points (means) and line segments (standard errors). Pure types are in filled symbols (blue for rural and orange for urban). Open symbols represent hybrid matings, with the color reflecting the maternal population origin.

## Discussion

Populations can adaptively respond to rapidly changing environments through evolutionary change and phenotypic plasticity occurring within or across generations or through their interaction (Merilä and Hendry 2014; Diamond and Martin 2016). Although distinguishing among these mechanisms is often difficult, doing so is necessary to understand and predict future responses to rapid environmental change (Urban et al. 2016). Here we used a multi-generational common-garden study to discriminate between the effects of transgenerational plasticity and the effects of evolutionary change. We explored this question in context of the adaptive divergence of thermal tolerance traits between urban and rural populations of the acorn ant *Temnothorax curvispinosus*. We found no evidence that transgenerational plasticity contributed to divergence between urban and rural populations in thermal tolerance, as shifts towards better heat tolerance and worse cold tolerance in urban populations were maintained over two generations in a common-garden laboratory environment. Further, we found that thermal tolerance traits were not inherited either maternally or paternally in the hybrid pairings.

Our results align with those of the few other studies that have used multi-generation common garden experiments in the context of thermal tolerance divergence to urban heat islands (Brans et al. 2017). Yet, the overall importance of transgenerational plasticity for evolutionary responses to novel environments is far from known. The thermal environment experienced by parents does appear to adaptively affect the thermal tolerance of their offspring in other contexts and systems, providing strong examples of anticipatory parental effects (Massamba-N’Siala et al. 2014; Chirgwin et al. 2018). However, a recent comprehensive synthesis of transgenerational plasticity across taxa and traits found relatively weak effects overall (Uller et al. 2013). Adaptive transgenerational plasticity in thermal tolerance could therefore prove to be relatively uncommon, albeit with important exceptions, such as those mentioned above. Alternatively, previous studies might not have been designed to detect such transgenerational plasticity effectively, or might not have focused on systems where such effects are expected to evolve (Burgess and Marshall 2014). This may occur if researchers are not actually manipulating the aspects of the environment that are most predictive of the conditions offspring will encounter (Uller 2008; Burgess and Marshall 2014).

Is there reason to expect adaptive parental effects to evolve in acorn ants as a response to the urban heat islands? Perhaps not. Acorn ant dispersal ability is quite limited (Herbers and Tucker 1986; Prebus 2017), and it is highly unlikely acorn ants could experience urban and rural temperature environments from one generation to the next. Further, within an acorn ant generation, workers experience substantial thermal variation over very rapid spatio-temporal scales, including diurnally within the acorn ant nest environment and spatially as foragers move throughout the landscape. This thermal variability is greater within urban forest patches, and urban population ants have evolved greater within-generation thermal plasticity to rapid temperature increases (Diamond et al. 2018b). Consequently, while both temperature means and variances differ between urban and rural populations, the temporal and spatial scale of this variation is unlikely to select for adaptive transgenerational effects.

Although the primary goal of our experiment was to evaluate transgenerational plasticity as an alternative explanation for the adaptive divergence in thermal tolerance between urban and rural *T. curvispinosus* populations, our study is not poised to address the overall presence or absence of parental effects. That is, we cannot rule out *any* role for transgenerational thermal plasticity in this system or others, only that it does not appear to explain the urban-rural divergence in thermal tolerance here. Indeed, because we reared acorn ant colonies under non-stressful temperatures within our multi-generation common garden experiment, it remains unclear whether other manipulations of the thermal environment, such as heat stress (Chirgwin et al. 2018) or seasonal temperature variation (Walsh et al. 2014), could induce transgenerational effects and whether such effects could have played an initial role in facilitating urban-rural divergence (Badyaev et al. 2002a,b; Badyaev 2008; Pfennig and Martin 2009, 2010).

While we did not find evidence for transgenerational plasticity as an explanation for the divergence in heat and cold tolerance traits among urban and rural acorn ant populations, the results of the urban-rural hybrid crosses yielded some surprising insights into the genetic architecture of acorn ant thermal tolerance. Specifically, our comparisons of heat and cold tolerance traits between the pure type and hybrid urban-rural crosses revealed that dominant and recessive alleles appear to underlie heat and cold tolerance traits, rather than being determined as purely additive traits. We found that both hybrid crosses resemble urban populations in heat tolerance and achieve greater cold tolerances similar to that of rural populations, rather than exhibiting intermediate phenotypes as would be expected for additive traits (Figure 1, Table 1). Why might both hybrid crosses express the greater heat tolerance of urban populations *and* the greater cold tolerance of rural populations? One explanation is that heat tolerance and cold tolerance are largely determined by different genes, with urban populations contributing a dominant allele(s) for heat tolerance and rural populations a dominant allele(s) for cold tolerance (Chown and Nicolson 2004). Alternatively, heat and cold tolerance could share genes with alternative conditional mutations (Griffiths et al. 2005), resulting in environmentally-dependent dominance. For example, imagine that urban and rural populations are fixed for different alleles of a heat shock protein. At low temperatures, the urban allele suffers loss of function and conversely the rural allele loses function at high temperatures, resulting in temperature-sensitive dominance. The latter scenario would suggest that the losses of cold tolerance in urban population could in part, be a correlated response to selection for increased heat tolerance in urban habitats. However, heat and cold tolerance are not strongly genetically correlated within colonies (Diamond et al. 2018a) suggesting that they may indeed be determined by separate genes. This would also correspond to the general findings from *Drosophila* and other ectotherm systems which show that cold and heat tolerance traits are under independent or semi-independent genetic control (Hoffmann et al. 2003).

Our research provides a case study in how an urban-rural cline in temperature can be used to explore the mechanisms underlying shifts in thermal tolerance trait values, specifically disentangling evolutionary change from transgenerational plasticity, and to understand the genetic architecture underlying these traits. But what else can we gain from such comparisons? With the growing consensus that evolution can occur in the wild on the scale of human lifetimes (Reznick et al. 2018), cities might also allow us to peer into the early stages of speciation as differences accumulate among urban and rural populations (Thompson et al. 2018). Indeed, as *Temnothorax curvispinosus* populations have evolved divergent thermal tolerances in response to urban heat islands, could this ecologically divergent selection promote speciation as well? Ecologically divergent environments can promote reproductive isolation in several ways. For example populations may become reproductively isolated by shifting the timing of reproduction between them (Rundle and Nosil 2005). In the acorn ant system, the date of peak alate production is shifted earlier by as much as 30 days in urban populations compared with rural populations, likely due to the differences in the timing of seasonal temperature cues in rural and urban habitats (Chick et al. 2019). This phenological shift in reproduction is thus likely to reduce possibilities for mating between urban and rural populations. In contrast, our laboratory breeding experiment shows that, given the opportunity, urban and rural alates (*i.e.*, reproductive-caste offspring) will mate and produce fertile offspring, revealing a lack of both behavioral prezygotic and intrinsic genetic postzygotic isolating mechanisms. Although, because male alates are haploid, any male sterility corresponding with Haldane’s rule would only be expressed by F2 male hybrids who through recombination, carry alleles from both urban and rural populations (Koevoets and Beukeboom 2009), which remains to be tested.

Ecologically divergent environments can also impose reproductive isolation by selecting against migrants and/or hybrid offspring (Nosil et al. 2003; Nosil 2004). Local adaptation and divergent selection between urban and rural thermal environments suggest that acorn ant migrants would face negative selection pressures in their non-native habitat. Hybrid acorn ant offspring, in comparison, inherit the combined thermal tolerance of both their parents, with heat and cold tolerances equaling those of urban and rural populations respectively (Figure 1, Table 1). This expanded thermal performance (Figure 2, Table 1) breaks the expectation of intermediate hybrid fitness for ecological speciation (Rundle and Nosil 2005) and suggests that hybridization could lead to population collapse rather than population divergence. We then expect that limited dispersal ability, shifted phenologies, or strong selection against migrants has enabled the evolutionary divergence between urban and rural populations in light of the expanded thermal breadth of their hybrid offspring. However, the evidence for any ongoing speciation between urban and rural population is both preliminary and mixed at this time.

In summary, the results from a two-generation common garden rearing experiment support the conclusion that acorn ants have evolved divergent heat and cold thermal tolerances in response to the selective agent of the urban heat island in Cleveland, Ohio. Surprisingly, thermal tolerance appears to be influenced by dominance rather than being a purely additive trait, and heat and cold tolerance might be determined by separate genes. While urban and rural populations have also diverged in their reproductive phenology, there is no evidence for other pre- or postzygotic reproductive isolation between them. *Temnothorax curvispinosus* populations have also diverged in thermal tolerance along at least one other urbanization gradient (Knoxville, Tennessee), and it is an open question remaining to be explored whether the genetic architecture, mating behavior or hybrid success is convergent across this independent selective gradient. As urban environments provide replicated theaters against which the evolutionary play unfolds, future work aimed at comparisons of the mechanisms of trait evolution across multiple urban heat islands could be especially fruitful.

## Acknowledgements

The Squire Valleevue and Valley Ridge Farm and the Holden Forests and Gardens provided access to field sites. M. Moore provided helpful comments. An Oglebay Fund grant provided financial support.

## Authors’ contributions

R.A.M. and S.E.D. designed the study, analyzed the data and wrote the first draft of the manuscript. All authors collected data and contributed to revisions.

## Data Accessibility Statement

Data will be deposited to Dryad upon acceptance.

## Supporting Information

**Table S1.** Geographic coordinates and percent developed impervious surface area (ISA) values (0% ISA indicates rural sites, 40-50% ISA indicates urban sites) of acorn ant collection sites.

| Site name | Longitude | Latitude | Source environment | ISA |
| --- | --- | --- | --- | --- |
| University Farm | -81.4245 | 41.49842 | Rural | 0 |
| Holden Arboretum | -81.3127 | 41.6088 | Rural | 0 |
| Ambler Park | -81.6055 | 41.4976 | Urban | 40 |
| Case Western Reserve University | -81.6137 | 41.50897 | Urban | 49 |
| Forest Hills | -81.5737 | 41.52751 | Urban | 44 |

**Table S2.**
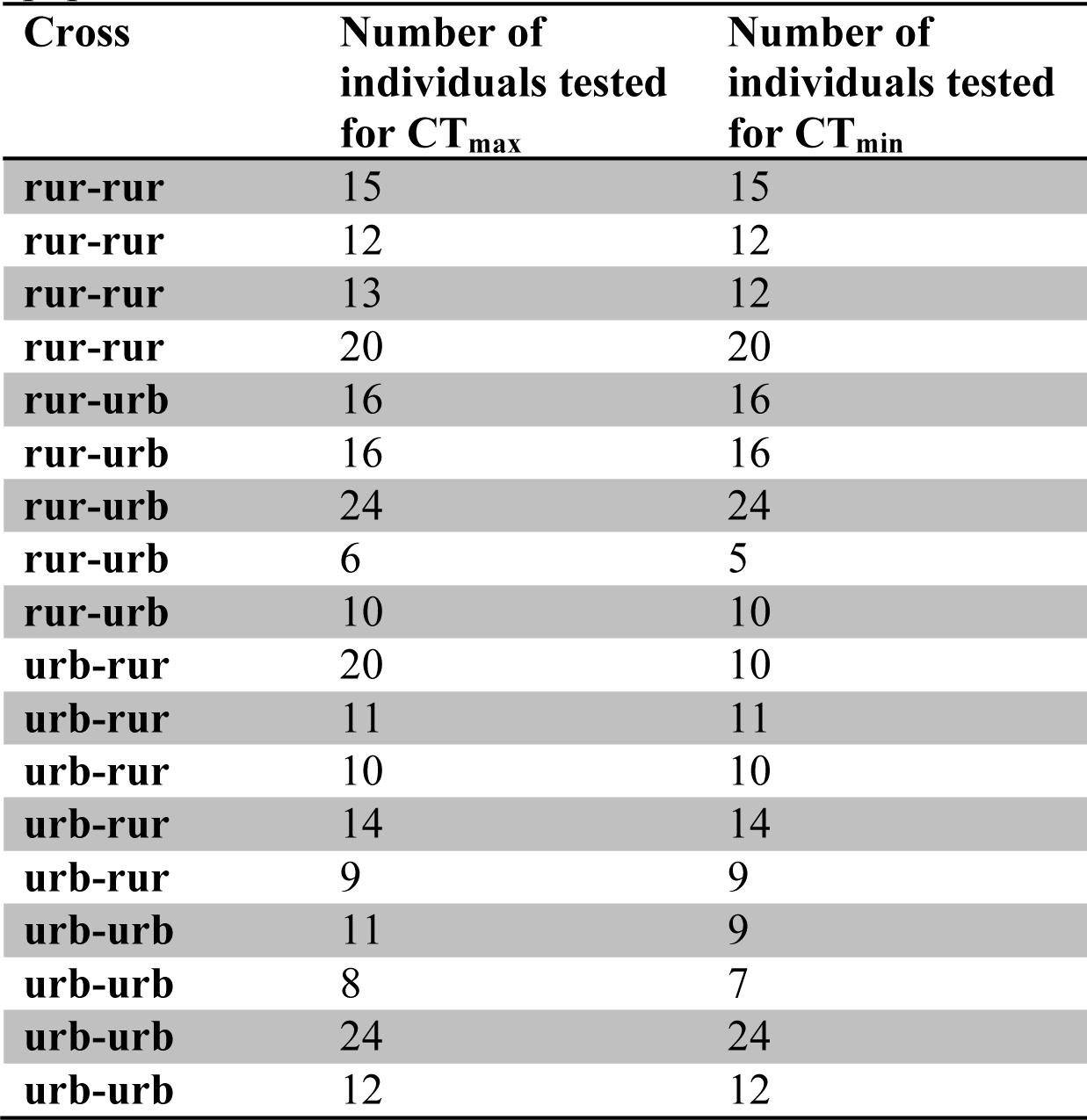
Number of individual worker ants tested for CT_min_ and CT_max_ per each colony. The type of cross (the two hybrids and two pure types) is also indicated with the maternal source population listed first.

| Cross | Number of individuals tested for $CT_{max}$ | Number of individuals tested for $CT_{min}$ |
| --- | --- | --- |
| rur-rur | 15 | 15 |
| rur-rur | 12 | 12 |
| rur-rur | 13 | 12 |
| rur-rur | 20 | 20 |
| rur-urb | 16 | 16 |
| rur-urb | 16 | 16 |
| rur-urb | 24 | 24 |
| rur-urb | 6 | 5 |
| rur-urb | 10 | 10 |
| urb-rur | 20 | 10 |
| urb-rur | 11 | 11 |
| urb-rur | 10 | 10 |
| urb-rur | 14 | 14 |
| urb-rur | 9 | 9 |
| urb-urb | 11 | 9 |
| urb-urb | 8 | 7 |
| urb-urb | 24 | 24 |
| urb-urb | 12 | 12 |

## References

Badyaev, A., G. Hill, and L. Whittingham. 2002a. Population consequences of maternal effects: sex-bias in egg-laying order facilitates divergence in sexual dimorphism between bird populations. Journal of Evolutionary Biology 15:997–1003.

Badyaev, A. V. 2008. Maternal effects as generators of evolutionary change. Annals of the New York Academy of Sciences 1133:151–161.

Badyaev, A. V., G. E. Hill, M. L. Beck, A. A. Dervan, R. A. Duckworth, K. J. McGraw, P. M. Nolan, and L. A. Whittingham. 2002b. Sex-biased hatching order and adaptive population divergence in a passerine bird. Science 295:316–318.

Bernardo, J. 1996. Maternal Effects in Animal Ecology. Integr Comp Biol 36:83–105.

Brans, K. I., M. Jansen, J. Vanoverbeke, N. Tüzün, R. Stoks, and L. D. Meester. 2017. The heat is on: Genetic adaptation to urbanization mediated by thermal tolerance and body size. Global Change Biology 23:5218–5227.

Burgess, S. C., and D. J. Marshall. 2014. Adaptive parental effects: the importance of estimating environmental predictability and offspring fitness appropriately. Oikos 123:769–776.

Chick, L. D., S. A. Strickler, A. Perez, R. A. Martin, and S. E. Diamond. 2019. Urban heat islands advance the timing of reproduction in a social insect. Journal of Thermal Biology 80:119–125.

Chirgwin, E., D. J. Marshall, C. M. Sgrò, and K. Monro. 2018. How does parental environment influence the potential for adaptation to global change? Proceedings of the Royal Society B: Biological Sciences 285:20181374.

Chown, S. L., and S. Nicolson. 2004. Insect Physiological Ecology: Mechanisms and Patterns. OUP Oxford.

Corl, A., K. Bi, C. Luke, A. S. Challa, A. J. Stern, B. Sinervo, and R. Nielsen. 2018. The genetic basis of adaptation following plastic changes in coloration in a novel environment. Current Biology 28:2970–2977.

DeWitt, T. J., and S. M. Scheiner. 2004. Phenotypic plasticity: functional and conceptual approaches. Oxford University Press.

Diamond, S. E., L. D. Chick, A. Perez, S. A. Strickler, and R. A. Martin. 2018a. Evolution of thermal tolerance and its fitness consequences: parallel and non-parallel responses to urban heat islands across three cities. Proc. Biol. Sci. 285.

Diamond, S. E., L. D. Chick, A. Perez, S. A. Strickler, and C. Zhao. 2018b. Evolution of plasticity in the city: urban acorn ants can better tolerate more rapid increases in environmental temperature. Conserv Physiol 6.

Diamond, S. E., L. Chick, A. Perez, S. A. Strickler, and R. A. Martin. 2017. Rapid evolution of ant thermal tolerance across an urban-rural temperature cline. Biol J Linn Soc 121:248–257.

Diamond, S. E., and R. A. Martin. 2016. The interplay between plasticity and evolution in response to human-induced environmental change. F1000Research 5.

Diamond, S. E., C. A. Penick, S. L. Pelini, A. M. Ellison, N. J. Gotelli, N. J. Sanders, and R. R. Dunn. 2013. Using Physiology to Predict the Responses of Ants to Climatic Warming. Integr Comp Biol 53:965–974.

Donelson, J. M., S. Salinas, P. L. Munday, and L. N. S. Shama. 2018. Transgenerational plasticity and climate change experiments: Where do we go from here? Global Change Biology 24:13–34.

Donihue, C. M., and M. R. Lambert. 2015. Adaptive evolution in urban ecosystems. Ambio 44:194–203.

Ghalambor, C. K., J. K. McKAY, S. P. Carroll, and D. N. Reznick. 2007. Adaptive versus non-adaptive phenotypic plasticity and the potential for contemporary adaptation in new environments. Functional Ecology 21:394–407.

Gorton, A. J., D. A. Moeller, and P. Tiffin. 2018. Little plant, big city: a test of adaptation to urban environments in common ragweed (Ambrosia artemisiifolia). Proceedings of the Royal Society B: Biological Sciences 285:20180968.

Griffiths, A. J., S. R. Wessler, R. C. Lewontin, W. M. Gelbart, D. T. Suzuki, J. H. Miller, and others. 2005. An introduction to genetic analysis. Macmillan.

Herbers, J. M. 1990. Reproductive Investment and Allocation Ratios for the Ant Leptothorax longispinosus: Sorting Out the Variation. The American Naturalist 136:178–208.

Herbers, J. M., and C. W. Tucker. 1986. Population Fluidity in Leptothorax Longispinosus (Hymenoptera: Formicidae).

Hoffmann, A. A., J. G. Sørensen, and V. Loeschcke. 2003. Adaptation of Drosophila to temperature extremes: bringing together quantitative and molecular approaches. Journal of Thermal Biology 28:175–216.

Imhoff, M. L., P. Zhang, R. E. Wolfe, and L. Bounoua. 2010. Remote sensing of the urban heat island effect across biomes in the continental USA. Remote Sensing of Environment 114:504–513.

Johnson, M. T. J., and J. Munshi-South. 2017. Evolution of life in urban environments. Science 358.

Kelly, M. 2019. Adaptation to climate change through genetic accommodation and assimilation of plastic phenotypes. Philosophical Transactions of the Royal Society B 374:20180176.

Koevoets, T., and L. W. Beukeboom. 2009. Genetics of postzygotic isolation and Haldane’s rule in haplodiploids. Heredity 102:16–23.

Lenth, R. 2018. emmeans: Estimated Marginal Means, aka Least-Squares Means.

Levis, N. A., A. J. Isdaner, and D. W. Pfennig. 2018. Morphological novelty emerges from pre-existing phenotypic plasticity. Nature ecology & evolution 2:1289.

Levis, N. A., and D. W. Pfennig. 2016. Evaluating “plasticity-first” evolution in nature: key criteria and empirical approaches. Trends in ecology & evolution 31:563–574.

Marshall, D. J., and T. Uller. 2007. When is a maternal effect adaptive? Oikos 116:1957–1963.

Massamba-N’Siala, G., D. Prevedelli, and R. Simonini. 2014. Trans-generational plasticity in physiological thermal tolerance is modulated by maternal pre-reproductive environment in the polychaete Ophryotrocha labronica. Journal of Experimental Biology 217:2004–2012.

Merilä, J., and A. P. Hendry. 2014. Climate change, adaptation, and phenotypic plasticity: the problem and the evidence. Evolutionary Applications 7:1–14.

Mousseau, T. A., and C. W. Fox. 1998. The adaptive significance of maternal effects. Trends Ecol. Evol. (Amst.) 13:403–407.

Nosil, P. 2004. Reproductive isolation caused by visual predation on migrants between divergent environments. Proceedings of the Royal Society of London. Series B: Biological Sciences 271:1521–1528.

Nosil, P., B. J. Crespi, and C. P. Sandoval. 2003. Reproductive isolation driven by the combined effects of ecological adaptation and reinforcement. Proceedings of the Royal Society of London. Series B: Biological Sciences 270:1911–1918.

Pfennig, D. W., and R. A. Martin. 2009. A maternal effect mediates rapid population divergence and character displacement in spadefoot toads. Evolution: International Journal of Organic Evolution 63:898–909.

Pfennig, D. W., and R. A. Martin. 2010. Evolution of character displacement in spadefoot toads: different proximate mechanisms in different species. Evolution: International Journal of Organic Evolution 64:2331–2341.

Pinheiro, J., D. Bates, S. DebRoy, D. Sarkar, and R Core Team. 2018. nlme: Linear and Nonlinear Mixed Effects Models.

Prebus, M. 2017. Insights into the evolution, biogeography and natural history of the acorn ants, genus Temnothorax Mayr (hymenoptera: Formicidae). BMC Evolutionary Biology 17:250.

Price, T. D.,A. Qvarnström, and D. E. Irwin. 2003. The role of phenotypic plasticity in driving genetic evolution. Proceedings of the Royal Society of London. Series B: Biological Sciences 270:1433–1440.

R Core Team. 2018. R: A Language and Environment for Statistical Computing. R Foundation for Statistical Computing, Vienna, Austria.

Reznick, D. N., J. Losos, and J. Travis. 2018. From low to high gear: there has been a paradigm shift in our understanding of evolution. Ecol. Lett., doi:10.1111/ele.13189.

Rivkin, L. R., J. S. Santangelo, M. Alberti, M. F. J. Aronson, C. W. de Keyzer, S. E. Diamond, M.-J. Fortin, L. J. Frazee, A. J. Gorton, A. P. Hendry, Y. Liu, J. B. Losos, J. S. MacIvor, R. A. Martin, M. J. McDonnell, L. S. Miles, J. Munshi□South, R. W. Ness, A. E. M. Newman, M. R. Stothart, P. Theodorou, K. A. Thompson, B. C. Verrelli, A. Whitehead, K. M. Winchell, and M. T. J. Johnson. 2019. A roadmap for urban evolutionary ecology. Evolutionary Applicatio 12:384–398.

Rundle, H. D., and P. Nosil. 2005. Ecological speciation. Ecology Letters 8:336–352.

Stuart, R. J., L. Gresham-Bissett, and T. M. Alloway. 1993. Queen adoption in the polygynous and polydomous ant, Leptothorax curvispinosus. Behav Ecol 4:276–281.

Terblanche, J. S., A. A. Hoffmann, K. A. Mitchell, L. Rako, P. C. le Roux, and S. L. Chown. 2011. Ecologically relevant measures of tolerance to potentially lethal temperatures. Journal of Experimental Biology 214:3713–3725.

Thompson, K. A., L. H. Rieseberg, and D. Schluter. 2018. Speciation and the City. Trends in Ecology & Evolution 33:815–826.

Uller, T. 2008. Developmental plasticity and the evolution of parental effects. Trends Ecol. Evol. (Amst>.) 23:432–438.

Uller, T., S. Nakagawa, and S. English. 2013. Weak evidence for anticipatory parental effects in plants and animals. J. Evol. Biol. 26:2161–2170.

Urban, M. C., G. Bocedi, A. P. Hendry, J.-B. Mihoub, G. Pe’er, A. Singer, J. R. Bridle, L. G. Crozier, L. D. Meester, W. Godsoe, A. Gonzalez, J. J. Hellmann, R. D. Holt, A. Huth, K. Johst, C. B. Krug, P. W. Leadley, S. C. F. Palmer, J. H. Pantel, A. Schmitz, P. A. Zollner, and J. M. J. Travis. 2016. Improving the forecast for biodiversity under climate change. Science 353:aad8466.

Walsh, M. R., D. Whittington, and C. Funkhouser. 2014. Thermal Transgenerational Plasticity in Natural Populations of Daphnia. Integr Comp Biol 54:822–829.

West-Eberhard, M. J. 2003. Developmental plasticity and evolution. Oxford University Press.

Winchell, K. M., R. G. Reynolds, S. R. Prado□Irwin, A. R. Puente□Rolón, and L. J. Revell. 2016. Phenotypic shifts in urban areas in the tropical lizard Anolis cristatellus. Evolution 70:1009–1022.

